# High Resolution Mitochondrial DNA Analysis Sheds Light on Human Diversity, Cultural Interactions and Population Mobility in Northwestern Amazonia

**DOI:** 10.1101/160218

**Authors:** Leonardo Arias, Chiara Barbieri, Guillermo Barreto, Mark Stoneking, Brigitte Pakendorf

**Author notes:** Leonardo Arias, Max-Planck-Institute for Evolutionary Anthropology, Deutscher Platz 6, D-04103, Leipzig, Germany. Grant sponsorship: Max Planck Society and COLCIENCIAS.

## Abstract

**Objectives:** Northwestern Amazonia (NWA) is a center of high linguistic and cultural diversity. Several language families and linguistic isolates occur in this region, as well as different subsistence patterns: some groups are foragers while others are agriculturalists. In addition, speakers of Eastern Tukanoan languages are known for practicing linguistic exogamy, a marriage system in which partners must come from different language groups. In this study, we use high resolution mitochondrial DNA sequencing to investigate the impact of this linguistic and cultural diversity on the genetic relationships and structure of NWA groups.

**Methods:** We collected saliva samples from individuals representing 40 different NWA ethnolinguistic groups and sequenced 439 complete mitochondrial genomes to an average coverage of 1030x.

**Results:** The mtDNA data revealed that NWA populations have high genetic diversity with extensive sharing of haplotypes among groups. Moreover, groups who practice linguistic exogamy have higher mtDNA diversity, while the foraging Nukak have lower diversity. We also find that rivers play a more important role than either geography or language affiliation in structuring the genetic relationships of populations.

**Discussion:** Contrary to the view of NWA as a pristine area inhabited by small human populations living in isolation, our data support a view of high diversity and contact among different ethnolinguistic groups; movement along rivers has probably facilitated this contact. Additionally, we provide evidence for the impact of cultural practices, such as linguistic exogamy, on patterns of genetic variation. Overall, this study provides new data and insights into a remote and little-studied region of the world.

---

Northwestern Amazonia (NWA) contains tremendous biological, linguistic, and cultural diversity, which likely reflects the heterogeneity of the landscape, especially the complex and extensive network of rivers found in this area. The region (Figure 1) extends from the Andean foothills in the west to the area between the Orinoco River and the Rio Negro in the east, and extends south until the confluence between the Rio Negro and the Amazon River. The northern border is defined by the Eastern Andean Cordillera and the Colombian-Venezuelan llanos, and in the south by the full length of the Putumayo River (Eriksen 2011).

In terms of linguistic diversity, NWA harbors ethnolinguistic groups belonging to the main South American language families accepted by specialists (Campbell 1997; Chacon 2014; Dixon and Aikhenvald 1999), namely Arawakan, Carib, Tupi, and Quechua. Additionally, several local families are also present, such as Tukanoan, Guahiban, Huitotoan, Boran, Peba-Yaguan, Piaroa-Saliban and Maku-Puinave, as well as various isolate languages like Tikuna, Cofan, and Kamentsa (Landaburu 2000). Furthermore, several indigenous groups live in voluntary isolation, and almost nothing is known about their linguistic affiliation (Franco 2002). The area has been proposed as the place of origin of the Arawakan family, since it contains the highest linguistic diversity within the family (Aikhenvald 1999; Heckenberger 2002; Zucchi 2002). In addition, all 20 languages of the Tukanoan family are found in the area; these are classified into two branches: The Western Tukanoan branch (WT) distributed along the Putumayo, Caquetá, and Napo rivers, and the Eastern Tukanoan branch (ET) along the Vaupés, Rio Negro, and Apaporis rivers and their tributaries (Chacon 2014). The language families Carib, Tupi, and Quechua are probably recent immigrants in NWA, since only one language per family is present in the area. In addition, the Tupi language Nheengatú or Lingua Geral is found in the region; however, this is a very recent introduction spread by missionaries during the 17^th^ and 18^th^centuries and by traders during the rubber boom in the 19^th^century, when it was used as a trade language (Sorensen 1967; Stenzel 2005).

**Figure 1.**
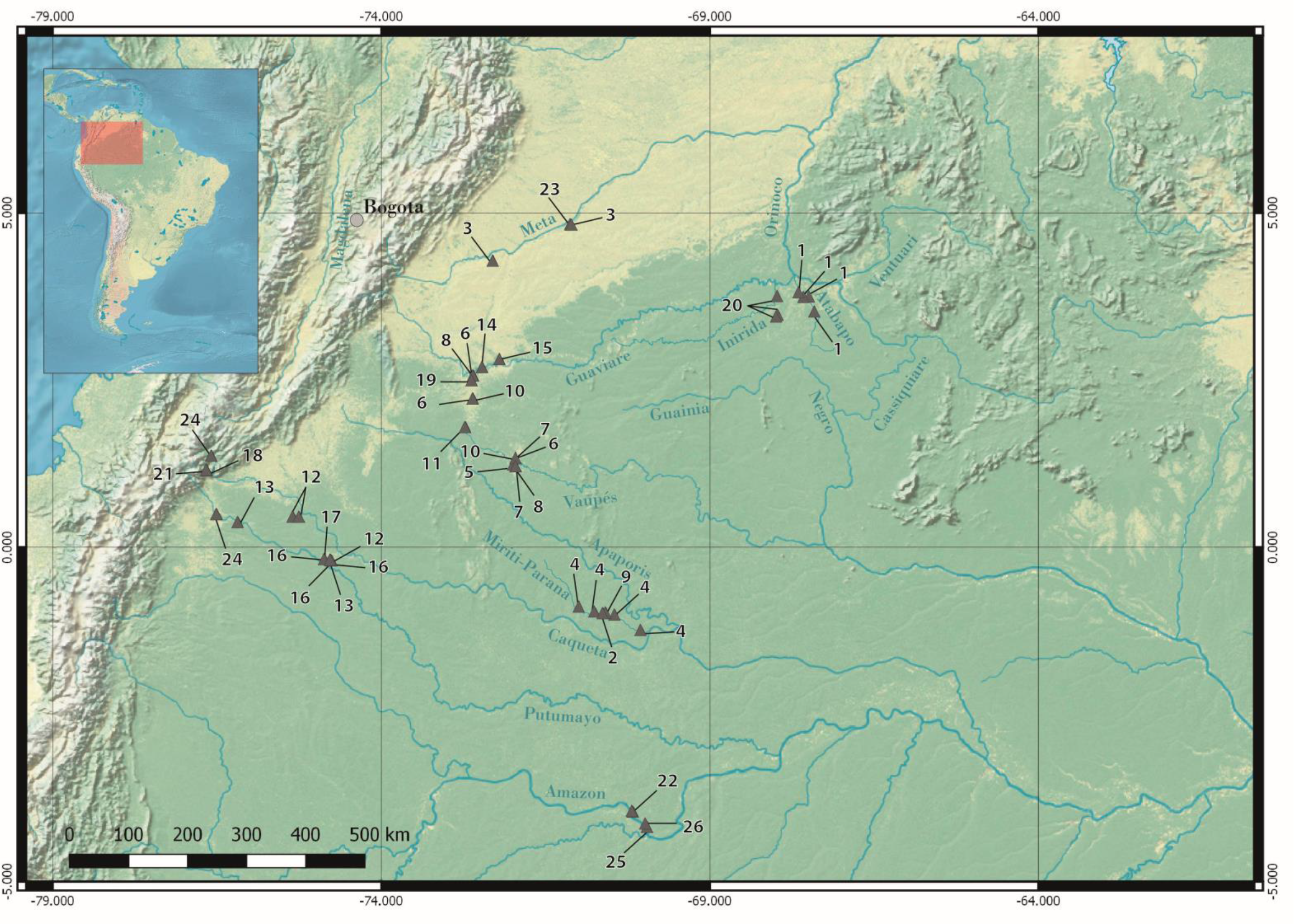
Geographic location of the sampling sites. Every triangle corresponds to a single community, which may contain more than one ethnolinguistic group. 1. Curripaco and Bare, 2. Matapi, 3. Ach-Piapoco, 4. Yucuna, 5. Carijona, 6. Desano, Yuruti, Pisamira, and Karapana, 7. Pira-Wanano, 8. Siriano, 9. Tanimuka, 10. Tukano, 11. Tuyuca and Tatuyo, 12. Coreguaje, 13. Siona, 14. Guayabero, 15. Sikuani, 16. Murui, 17.Uitoto, 18. Kamentsa, 19. Nukak, 20. Puinave, 21. Pasto, 22. Yagua, 23. Saliba, 24. Inga, 25. Tikuna, 26. Cocama.

In terms of cultural diversity, while NWA has often been viewed as a pristine area inhabited only by small, isolated, seminomadic tribes with an economy based on hunting and gathering (Denevan 1992; Meggers 1954), in fact there is considerable variation in subsistence and marriage practices. While some groups are traditional foragers, others engage in agriculture, and instead of isolated groups, archaeological and anthropological evidence now shows that NWA was indeed part of a continent-wide network of exchange and trade. Complex societies organized in chiefdoms and multiethnic confederations arose in the region, and multilingualism and extensive interactions among ethnolinguistic groups were the norm (Heckenberger 2002; Hornborg 2005; Santos-Granero 2002; Vidal 1997). In particular, the groups speaking Eastern Tukanoan languages and some of their Arawakan neighbors living in the basin of the Vaupés River and Rio Negro engage in an exceptional marital practice known as linguistic exogamy (Aikhenvald 1996; Chacon and Cayón 2013; Sorensen 1967; Stenzel 2005). According to this cultural norm, individuals are required to marry someone from a different language group, with each individual’s linguistic affiliation determined by the language of the father. Linguistic exogamy thus creates a situation of multilingualism and movement of people (especially women, since it is accompanied by patrilocality and patrilineality) among the groups participating in the system (Sorensen 1967).

Historical linguistics, cultural anthropology, and archaeology are the main disciplines that have traditionally addressed questions regarding the origins, prehistory, and genetic relationships among the NWA ethnolinguistic groups (Campbell 1997; Chacon 2014; Heckenberger 2008; Lathrap 1970; Meggers 1948; Nettle 1999). However, due to the scarcity of the archaeological record, the time-depth limitations of linguistic methods based on lexical cognates to establish deep relationships (Dediu and Levinson 2012; Hock and Joseph 2009), and the insufficiency of documentation and description of a large number of the NWA societies, many of these questions remain to be fully answered. The oldest archaeological evidence of human occupation in NWA comes from a single site on the middle Caquetá River dated between 9250 and 8100 BP, containing a great variety of stone artifacts, carbonized seeds and other botanical remains from different palm species, as well as phytoliths of bottle gourd, leren, and pumpkin (Aceituno et al. 2013; Gnecco and Mora 1997), indicating that these early human groups relied on vegetable resources that are still exploited by contemporary societies in NWA.

One hypothesis about the peopling of NWA was proposed by Nimuendajú (1950), who suggested that the region was first inhabited by hunter-gatherer populations, perhaps the ancestors of the Maku-Puinave groups, most of whom still practice a foraging lifestyle. Proto-Arawakan groups then started expanding in the region from their place of origin located between the Orinoco river and the Rio Negro (Heckenberger 2002; Lathrap 1970), and finally, the Tukanoans are assumed to have arrived in the area and displaced peoples speaking Arawakan and Maku-Puinave languages from the Vaupés (the Tukanoans probably came from the Napo-Putumayo, where Western Tukanoans still live). However, this scenario does not account for the presence of groups belonging to the Carib, Guahiban, Huitotoan, and Boran language families and the various language isolates in the region.

Genetic studies can provide insights into population history, and indeed studies of mitochondrial DNA (mtDNA) genetic variation in Native American populations have contributed greatly to our knowledge about the peopling of the Americas. Early studies using restriction fragment length polymorphisms (RFLP) and sequencing of the hypervariable region one (HVS-I) identified five founder lineages or haplogroups, designated as A to D and X (Bailliet et al. 1994; Barbieri et al. 2011; Gaya-Vidal et al. 2011; Keyeux et al. 2002; Lewis et al. 2007; Schurr 2004; Torroni et al. 1993). Whereas haplogroups A-D are widely distributed in the Americas, haplogroup X is restricted to North America (Bolnick and Smith 2003; Malhi et al. 2001). The analysis of HVS-I in several Native American populations showed that haplogroups A-D exhibit similar levels of diversity (Bonatto and Salzano 1997), supporting the hypothesis of a single origin of all Native American populations from a Northeast Asian source. Additionally, HVS-I data have been used to determine the genetic relationships among indigenous populations in South America and to test hypotheses concerning how genetic variation is structured at the regional and continental levels (Barbieri et al. 2011; Gaya-Vidal et al. 2011; Lewis et al. 2007; Marrero et al. 2007; Melton et al. 2007). These studies revealed that Andean (or western) populations show higher levels of diversity and low genetic distances in contrast to the Eastern populations, who show the opposite pattern. However, in previous studies NWA populations have been generally underrepresented, and hence the inferences about the genetic structure of the entire Amazonian region are based on a small number of populations.

Recent developments in sequencing technology allow the determination of complete mtDNA genomes at the population level and thus enable unbiased insights into the maternal history of human populations (Delfin et al. 2014; Gunnarsdottir et al. 2011; Kivisild 2015). At present, no such studies are reported for South American indigenous populations. Available studies of complete mtDNA genomes from Native Americans have been restricted to a limited number of individuals carrying particular haplogroups, usually selected based on their HVS-I sequences (Achilli et al. 2013; Bodner et al. 2012; de Saint Pierre et al. 2012; Fagundes et al. 2008; Lee and Merriwether 2015; Perego et al. 2009; Perego et al. 2010), or to archaeological remains from different time periods (Fehren-Schmitz et al. 2015; Llamas et al. 2016). These studies have primarily focused on inferences about the peopling of the continent, the number of migrations, the divergence times, and changes in the effective population size through time. Nevertheless, several problems and biases are associated with this sampling strategy. First, the overall diversity might be underestimated, since individuals carrying the same HVS-I sequence can exhibit considerable variation in the coding region (Gunnarsdottir et al. 2011). Secondly, the reconstruction of demographic trends can be skewed, since the estimation of effective population sizes through time using Bayesian coalescent methods (i.e. Bayesian skyline plots in BEAST) can generate spurious signals of population growth when based on samples selected by haplogroup (Gunnarsdottir et al. 2011). Lastly, the histories and origins of specific populations cannot be investigated, since the coalescent age of a particular lineage does not correspond to the age of the population, especially when the diversity within each lineage is unknown (Schurr 2004). In this study, we use complete mtDNA sequencing in a large and representative sample of populations covering the extant ethnolinguistic diversity from NWA to reconstruct their maternal history, as well as to determine their genetic diversity and to make inferences about the origins of this diversity.

Finally, we aim to investigate the impact of prehistoric population dynamics and cultural interactions on the structure of the genetic variation observed among present-day NWA populations.

## MATERIALS AND METHODS

### Sample collection

Samples from unrelated individuals belonging to 40 ethnolinguistic groups were collected during several expeditions carried out by one of the authors (L.A.) in five departments (administrative divisions) of NWA, namely: Amazonas, Guainía, Guaviare, Meta and Putumayo (Table 1, Figure 1). The samples consisted of either saliva, collected as 3 mL of saliva in 3 mL of lysis buffer (Quinque et al. 2006), or blood samples stabilized with EDTA. Written informed consent was obtained from each participant, and from the community leader and/or local/regional indigenous organizations, after giving a full description of the aims of the study. Local translators and fieldwork assistants helped to explain and translate into the local languages when individuals or communities were not proficient in Spanish. Additionally, each participant answered a short questionnaire soliciting information regarding their birthplace, language, ethnic affiliation and that of their parents and grandparents. The study was approved by the ethics committee of the Universidad del Valle in Cali, Colombia and the Ethics Commission of the University of Leipzig Medical Faculty. All procedures were undertaken in accordance with the Declaration of Helsinki on ethical principles and an export permit was issued by the Colombian Ministry of Health and Social Protection.

**Table 1.** Sampled ethnolinguistic groups with information on merged groups (see Material & Methods: Data Analysis) given below the compound names.

| Population | Label in figure 1 | N | Census Sizea | Language family | Subsistence Strategyb | River/place of residence |
| --- | --- | --- | --- | --- | --- | --- |
| Yucu-Matapi |  | 39 |  | Arawakan | AG | Mirití-Paraná |
| Yucuna | 4 | 31 | 550 | Arawakan | AG | Mirití-Paraná |
| Matapi | 2 | 8 | 220 | Arawakan | AG | Mirití-Paraná |
| Curripaco |  | 17 |  | Arawakan | AG | Atabapo |
| Curripaco | 1 | 16 | 7827 | Arawakan | AG | Atabapo |
| Bare | 1 | 1 | NAC | Arawakan | AG | Atabapo |
| Ach-Piapoco |  | 24 |  | Arawakan | AG | Meta |
| Achagua | 3 | 6 | 283 | Arawakan | AG | Meta |
| Piapoco | 3 | 18 | 4926 | Arawakan | AG | Meta |
| *Cabiyarid* |  | 1 | 311 | Arawakan | AG | Mirití-Paraná |
| Carijona | 5 | 8 | 307 | Carib | AG | Upper-Vaupés |
| *Cofan* |  | 6 | 877 | Cofan | AG | Guamúez |
| *Barasano* |  | 4 | 2008 | Eastern Tukanoan | AG | Upper-Vaupés |
| Desano | 6 | 17 | 2457 | Eastern Tukanoan | AG | Upper-Vaupés |
| *Kubeo* |  | 5 | 6647 | Eastern Tukanoan | AG | Upper-Vaupés |
| Other-EET |  | 10 |  | Eastern Tukanoan | AG | Upper-Vaupés |
| Tuyuca | 11 | 7 | 642 | Eastern Tukanoan | AG | Upper-Vaupés |
| Yuruti | 6 | 1 | 687 | Eastern Tukanoan | AG | Upper-Vaupés |
| Pisamira | 6 | 1 | 61 | Eastern Tukanoan | AG | Upper-Vaupés |
| Karapana | 6 | 1 | 464 | Eastern Tukanoan | AG | Upper-Vaupés |
| Pira-Wanano |  | 13 |  | Eastern Tukanoan | AG | Upper-Vaupés |
| Piratapuyo | 7 | 8 | 697 | Eastern Tukanoan | AG | Upper-Vaupés |
| Wanano | 7 | 5 | 1395 | Eastern Tukanoan | AG | Upper-Vaupés |
| Siriano | 8 | 10 | 749 | Eastern Tukanoan | AG | Upper-Vaupés |
| Tanimuka | 9 | 10 | 1247 | Eastern Tukanoan | AG | Mirití-Paraná |
| Tuka-Tatuyo |  | 10 |  | Eastern Tukanoan | AG | Upper-Vaupés |
| Tukano | 10 | 8 | 6996 | Eastern Tukanoan | AG | Upper-Vaupés |
| Tatuyo | 11 | 2 | 331 | Eastern Tukanoan | AG | Upper-Vaupés |
| Siona | 13 | 17 | 734 | Western Tukanoan | AG | Putumayo |
| Coreguaje | 12 | 19 | 2212 | Western Tukanoan | AG | Caquetá |
| Sikuani | 15 | 16 | 23006 | Guahiban | HGP | Guaviare |
| Guayabero | 14 | 35 | 1118 | Guahiban | HGP | Guaviare |
| Saliba | 23 | 16 | 1929 | Piaroa-Saliban | AG | Meta |
| Mur-Uitotoe |  | 26 | 7343 | Huitotoan | AG | Putumayo |
| Murui | 16 | 18 |  | Huitotoan | AG | Putumayo |
| Uitoto | 17 | 8 |  | Huitotoan | AG | Putumayo |
| Puinave | 20 | 19 | 6604 | Maku-Puinave | HGP | Inirida |
| Nukak | 19 | 16 | 1483 | Maku-Puinave | HGP | Interfluvial |
| Pasto | 21 | 14 | 69789 | Pasto | AG | Andean |
| Kamentsa | 18 | 11 | 4773 | Kamentsa | AG | Andean |
| Inga | 24 | 17 | 19079 | Quechuan | AG | Andean |
| Tikuna | 25 | 18 | 7102 | Tikuna | AG | Amazonas |
| Cocama | 26 | 17 | 792 | Tupi | AG | Amazonas |
| Yagua | 22 | 13 | 297 | Peba-Yaguan | AG | Amazonas |
| *Guambiano* |  | 1 | 23462 | Barbacoan | AG | Andean |
| *Nasa* |  | 1 | 138501 | Nasa | AG | Andean |
| *Mestizo* |  | 9 | NA | Mestizo | NA | NA |
| Total |  | 439 |  |  |  |  |

a. Data from: (Arango and Sánchez 2004). b. AG: agriculturalist; HGP: Hunter-gatherer populations, data from D-PLACE (Kirby et al 2016).and HG (https://huntergatherer.la.utexas.edu/home, accessed on 06.06.2017). d. Populations with label in italics were not considered in the population-based analyses. e. Census data reports the population size including groups that speak five dialectal varieties.

### DNA sequencing and sequence processing

The DNA was extracted from blood samples with the “salting out” method (Miller et al. 1988) and from the saliva samples with the QIAamp DNA Midi kit (Qiagen), starting from 2.0 mL of the saliva/buffer mixture. The concentration of DNA was quantified with the NanoDrop 8000 spectrophotometer (Thermo Scientific). We prepared genomic libraries with double indices and enriched for full mtDNA genomes using a hybridization-capture method described previously (Kircher et al. 2012; Maricic et al. 2010). From the enriched libraries, paired-end sequences of 100 bp length were generated on the Illumina Hiseq 2500 platform. Base-calling was performed using freeIbis (Renaud et al. 2013), and Illumina adapters were trimmed and completely overlapping paired sequences were merged using leeHOM (Renaud et al. 2014a). The sequencing data were de-multiplexed using deML (Renaud et al. 2014b) and the sequences aligned against the human reference genome 19 using BWA’s *aln* algorithm (Li and Durbin 2009). After duplicate removal using PicardTools v2.1.1 (https://github.com/broadinstitute/picard). we performed an iterative alignment for each library individually to obtain mtDNA consensus sequences. In the first step, we extracted all sequencing reads of a library that aligned either to the mitochondrial genome or to a list of nuclear copies of mtDNA (NUMTs) (Li et al. 2012). We subsequently aligned these reads to the revised Cambridge Reference Sequence (rCRS; Andrews et al. 1999) using BowTie2’s *very-sensitive* algorithm (Langmead and Salzberg 2012) and called a consensus sequence. In the second step, the reads were re-aligned to the library’s respective consensus sequence generated in the first step, using the same BowTie2 settings. After the second alignment step, we called a final consensus sequence that was used throughout the rest of the analysis. Final sequences in fasta format were aligned to the Revised Cambridge Reference Sequence (rCRS (Andrews et al. 1999)) with the multiple sequence alignment software Mafft (Katoh and Standley 2013), and manually inspected for alignment errors with Bioedit ver. 7.2.5 (Hall 1999). The two poly-C regions (np 303-315 and 16.183-16,194) were excluded from the subsequent analysis; although one position (16,189) diagnostic for haplogroup B2 is therefore not considered in the haplogroup-calling analysis, the additional substitutions defining this haplogroup that occur elsewhere in the mitochondrial genome enable unambiguous assignment of sequences to lineage B2.

### Data analysis

We considered populations with a sample size of 10 individuals or more, and merged populations with sample sizes smaller than 10 based on linguistic criteria when our initial analyses did not show significant genetic differences, as follows (Table 1). The Arawakan groups Achagua (n=6) and Piapoco (n=18) were merged into a single population, since their indigenous reservations are adjacent and individuals often intermarry (data available on request); one Bare (n=1) individual was added to the Curripaco (n=16) sample among whom he was living when sampled on the Atabapo River; Yucuna (n=31) and Matapi (n=8) were merged into a single population, since they both speak Yucuna, live along the same river, and intermarry (data available on request); and Murui (n=18) and Uitoto (n=8) were merged, as these two groups belong to the same language family, which is composed of several dialects that are mutually intelligible (http://glottolog.org/resource/languoid/id/huit1251. accessed on 31.05.2017). Finally, following the latest classification of the Tukanoan family (Chacon 2014), the Eastern Tukanoan groups Piratapuyo (n=8) and Wanano (n=5) were merged as Pira-Wanano; Tukano (n=8) and Tatuyo (n=2) were merged as Tuka-Tatuyo; and Tuyuca (n=7), Yuruti (n=1), Pisamira (n=1), and Karapana (n=1) were merged as Other-ET. The only group with a sample size smaller than 10 that we retained as a separate group in the analyses were the Carijona (n=8), since this is the only Carib-speaking group living in NWA. Moreover, they are at risk of disappearing both physically and culturally, with less than 30 active speakers of Carijona scattered in two communities, and they occupy an important place in the ethno-history of the region (Franco 2002). We excluded Barasano (n=4), Kubeo (n=5), Cofan (n=6), Cabiyari (n=1), Guambiano (n=1), and Nasa (n=1) from all the analyses except the haplotype networks, since this analysis represents the evolutionary relationships among individual sequences. We furthermore excluded nine individuals with maternal ancestry outside of NWA (labeled ‘Mestizo’ in Table 1) from all analyses. After merging and filtering as described above, 412 sequences from 24 groups were kept in the population-based analyses.

Based on information from D-PLACE (Kirby et al. 2016) and HG database (https://huntergatherer.la.utexas.edu/home. accessed on 06.06.2017), we divided the populations into agriculturalists (AG) and hunter-gatherers (HGP). In the latter category we included the Nukak, who currently still practice a foraging way of life, as well as the Puinave, Sikuani, and Guayabero, who have all adopted agriculture only recently (Kondo 2002; Uribe Tobón and Instituto Colombiano de Cultura 1992)

The haplogroup affiliation of the individual sequences was determined with Haplogrep (Kloss-Brandstatter et al. 2011), based on Phylotree build 16 (van Oven and Kayser 2009). Haplogroup frequencies by population were estimated by simple counting, and a correspondence analysis (CA) based on the frequency of sub-haplogroups (e.g. A2a) was performed and visualized with the R-packages FactoMineR (Le et al. 2008) and factoextra (Kassambara and Mundt 2016), respectively.

Population-based statistical analyses were performed with Arlequin v3.5 (Excoffier and Lischer 2010); these include the analysis of molecular variance (AMOVA), estimation of molecular diversity indices, the estimation of pairwise genetic distances based distances based on Φ_ST_, and Tajima’s D test of selective neutrality. A Multidimensional Scaling analysis (MDS) was performed on the matrix of pairwise Φ_ST_ values to visualize the distances between populations. Additionally, we performed a Mantel test to evaluate the correlations between genetic distances and geographic distances. The matrix of geographic distances was built using the geographic coordinates of the location where the majority of samples for each ethnolinguistic group were collected and then calculating the great circle distances between locations via the R packages ade4 and geosphere (Dray and Dufour 2007; Hijmans 2016). Furthermore, a multiple regression analysis on distance matrices (MRM) (Goslee and Urban 2007) with the form: MRM(as.dist(gen.dist) ∼ as.dist(geo.dist) + as.dist(rivers.dist)) was performed; the analysis takes into consideration a matrix of geographic distances and a matrix of proximity along rivers as predictor variables of the genetic distances (pairwise Φ_ST_ values) between populations (Pugach et al. 2016; Yunusbayev et al. 2012). For the matrix of river distances a value of zero was given to populations living along the same river or on rivers that are closely connected, and a value of one was given to populations living on different rivers.

The sharing of haplotypes between populations was estimated with in-house R scripts as the proportion of pairs of identical sequences shared between populations. Additionally, networks of haplotypes were constructed with the software Network ver. 4.6.1.3 and visualized with Network Publisher ver. 2.0.0.1 (http://www.fluxus-engineering.com). Finally, Bayesian skyline plots (BSP) were constructed with BEAST ver. 1.8.2 (Drummond et al. 2012). For this analysis, the best substitution model was estimated with jModeltest 2.1.7 (Darriba et al. 2012), and BEAST was used to estimate whether a strict or a relaxed clock model best fits the data. This analysis was performed on both the complete sequences and the sequences partitioned into coding (577-16023) and non-coding (16024-576) regions, applying the corresponding substitution rates reported previously (Soares et al. 2009).

## RESULTS

We generated 439 complete mitochondrial sequences to an average coverage per sample of 1030x, which were deposited in GenBank with accession numbers: XXXXXXXX-XXXXXXXX and XXXXXXXX-XXXXXXXX. All sequences belonged to one of the main Native American haplogroups, namely A2, B2, C1 and D1. Haplogroups A2 and C1 were the most frequent lineages in the NWA populations (excluding the so-called ‘Mestizos’), with more than half of all sequences belonging to A2 (90 haplotypes in 138 sequences) and C1 (95 haplotypes in 181 sequences) together; in contrast, B2 (49 haplotypes in 73 sequences) and D1 (32 haplotypes in 38 sequences) were less frequent. Table 2 provides a breakdown of the haplogroup frequencies for the ethnolinguistic groups included in the population analyses.

**Table 2.** Frequency of haplogroups for the 24 NWA ethnolinguistic groups included in the population analyses

| Population | N | A2 | B2 | C1 | D1 |
| --- | --- | --- | --- | --- | --- |
| Yucu-Matapi | 39 | 0.28 | 0.10 | 0.56 | 0.05 |
| Curripaco | 17 | 0.18 | 0.53 | 0.24 | 0.06 |
| Ach-Piapoco | 24 | 0.54 | 0.04 | 0.42 | 0.00 |
| Carijona | 8 | 0.13 | 0.25 | 0.63 | 0.00 |
| Desano | 17 | 0.29 | 0.12 | 0.41 | 0.18 |
| Other-ET | 10 | 0.50 | 0.00 | 0.30 | 0.20 |
| Pira-Wanano | 13 | 0.31 | 0.08 | 0.38 | 0.23 |
| Siriano | 10 | 0.40 | 0.10 | 0.40 | 0.10 |
| Tanimuka | 10 | 0.50 | 0.10 | 0.40 | 0.00 |
| Tuka-Tatuyo | 10 | 0.30 | 0.10 | 0.30 | 0.30 |
| Siona | 17 | 0.59 | 0.35 | 0.00 | 0.06 |
| Coreguaje | 19 | 0.11 | 0.16 | 0.74 | 0.00 |
| Sikuani | 16 | 0.25 | 0.00 | 0.75 | 0.00 |
| Guayabero | 35 | 0.43 | 0.23 | 0.34 | 0.00 |
| Saliba | 16 | 0.19 | 0.06 | 0.56 | 0.19 |
| Mur-Uitoto | 26 | 0.23 | 0.12 | 0.31 | 0.35 |
| Puinave | 19 | 0.11 | 0.42 | 0.47 | 0.00 |
| Nukak | 16 | 0.00 | 0.31 | 0.69 | 0.00 |
| Pasto | 14 | 0.36 | 0.14 | 0.21 | 0.29 |
| Kamentsa | 11 | 0.18 | 0.09 | 0.64 | 0.09 |
| Inga | 17 | 0.59 | 0.06 | 0.29 | 0.06 |
| Tikuna | 18 | 0.44 | 0.00 | 0.44 | 0.11 |
| Cocama | 17 | 0.29 | 0.35 | 0.29 | 0.06 |
| Yagua | 13 | 0.62 | 0.15 | 0.23 | 0.00 |

A Correspondence Analysis (CA) (Figure 2) shows the clustering of populations based on the frequency of sub-haplogroups. We observed differences among populations without a clear clustering by language family, with the exception of the Eastern Tukanoan groups Siriano, Desano, Pira-Wanano, Tuka-Tatuyo and Other-ET, which were near one another in the left side of the plot. However, Tanimuka, who also speak an Eastern Tukanoan language, were far apart from their linguistic relatives. Additionally, Guayabero and Sikuani (who speak languages belonging to the Guahiban family) were located close to each other in the lower left pane of the plot. In addition to language affiliation, a few cases of proximity in the CA plot could be attributed to geographic proximity, as in the case of Kamentsa, Pasto and Inga, who all live close to one another in the Andean foothills. In other cases, the relatively close proximity of populations could be attributed to their being settled along the same river or on rivers that are part of the same basin (Supporting information Figure S1), as in the case of Curripaco and Puinave, who live on the Inírida and Atabapo rivers.

**Figure 2.**
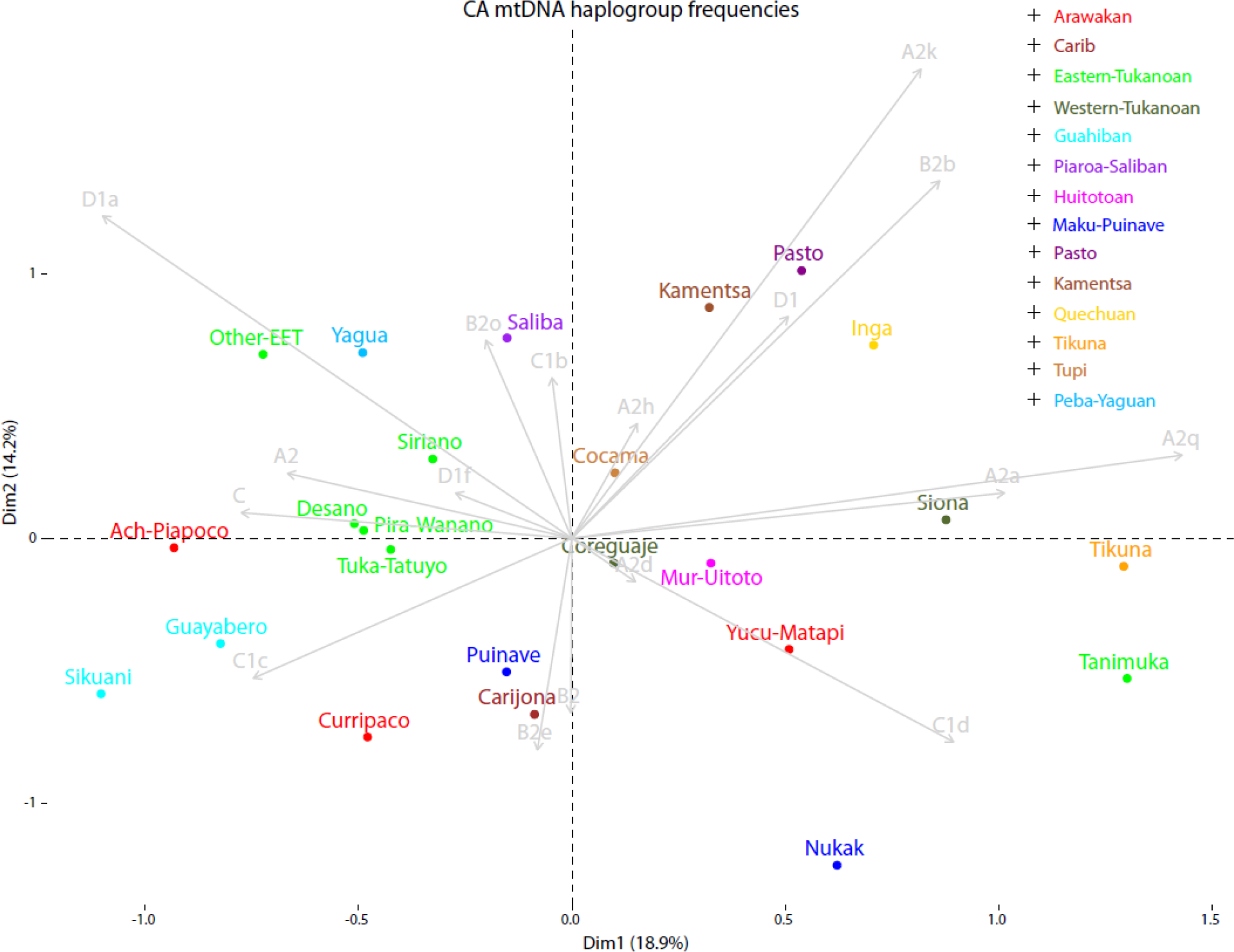
Correspondence analysis based on the sub-haplogroup frequencies by population. Populations are color-coded by linguistic affiliation.

### Molecular diversity indices

The genetic variation in these communities was assessed through different molecular diversity indices (Figure 3, Supporting information Table S1). On average, the gene diversity in these groups was high (0.9), but there were differences amongst them. For example, Eastern Tukanoan groups showed consistently high values of gene diversity, with the exception of the Tanimuka, who had one of the lowest values (0.73). The Western Tukanoan groups Coreguaje (0.92) and Siona (0.82) showed lower values than Eastern Tukanoan groups. Among Arawakans, the Ach-Piapoco had the lowest value (0.77). The hunter-gatherer group Nukak showed the lowest gene diversity of all groups (0.64): only four haplotypes were observed among the 16 individuals analyzed. Additionally, we observed that agriculturalist groups tended to have higher gene diversities (average = 0.92) than hunter-gatherer groups (average = 0.80) (Mann-Whitney U test, P-value = 0.03).

**Figure 3.**
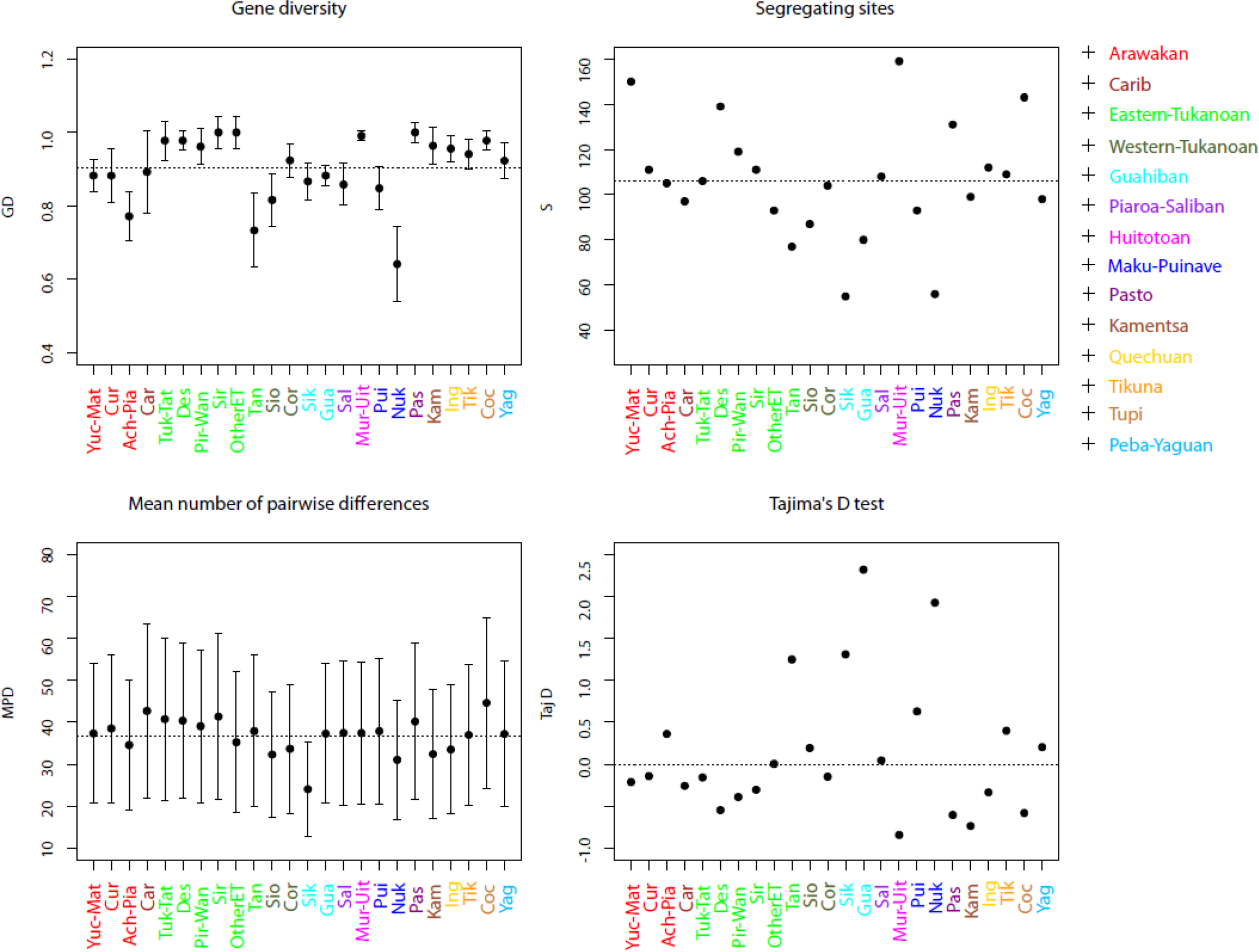
Molecular diversity indices by population. Dashed lines correspond to average values, except for Tajima’s D test which corresponds to zero. Populations are color-coded by linguistic affiliation as in Figure 2.

The mean number of pairwise differences (MPD) per population showed less variation, with an average of 41.07 +/- 17.86 differences. The smallest values were found in Sikuani (24.08 +/- 11.18) and Nukak (31.07 +/- 14.33), and the largest values were observed in Cocama (44.64 +/- 20.37), Carijona (42.71 +/- 20.8), and Siriano (41.38 +/- 19.66) The D values of Tajima’s test of neutrality (Tajima 1989) ranged from -0.735 to 2.318. Under neutrality, Tajima’s D is expected to be equal to zero and significant departures are interpreted as a result of selection or changes in population size. Although none of the D values were significant (all P-values > 0.2), positive D values >1.2 were obtained for Guayabero, Nukak, Sikuani and Tanimuka, which may reflect recent reductions in the size of these populations. This hypothesis was supported both by the distribution of pairwise differences by population (Supporting information Figure S2), which showed increased frequencies for the category of small differences (0 and 1 differences) and for the category of large differences (50 or more), as well as by the Bayesian reconstruction of population size changes through time (BSP plots, Supporting information Figure S3 and below). Furthermore, the Tanimuka and Nukak had the lowest gene diversity values.

### Shared haplotypes

A total of 216 different haplotypes were observed among the 412 sequences included in this analysis, pointing to a considerable number of shared sequences. Of these, 146 were unique haplotypes and 70 haplotypes were shared among 266 sequences: 52 within populations, 31 between populations, and 13 both within and between populations. The shared haplotypes accounted for 64.6% of all the sequences analyzed. This amount of haplotype sharing between populations is considerably high when compared to other population-based studies of complete mitochondrial genomes (Table 3). In other studies, the majority of shared haplotypes were generally observed within populations, with the exception of two African datasets from Burkina Faso and Zambia (Barbieri et al. 2013; Barbieri et al. 2012), which showed low levels of sharing both within and between populations. The highest level of sharing between populations was observed for Siberian populations spread over a large geographic area (Duggan et al. 2013); the NWA populations analyzed in this study showed the second highest value of sharing between populations.

**Table 3.** Shared haplotypes in a worldwide sample of complete mitochondrial sequences sampled at the population level

| Geographic region | #Sequences | #Haplotypes | %Unique haplotypes | Shared Within population | Shared Between populations | Source |
| --- | --- | --- | --- | --- | --- | --- |
| NW Amazonia | 412 | 216 | 0.676 | 0.241 | 0.144 | Present study |
| Burkina Faso | 335 | 332 | 0.991 | 0.006 | 0.003 | (Barbieri et al. 2012) |
| SW Zambia | 169 | 146 | 0.897 | 0.048 | 0.055 | (Barbieri et al. 2013) |
| Botswana/Namibia | 218 | 128 | 0.75 | 0.188 | 0.133 | (Barbieri et al. 2014) |
| Philippines | 365 | 233 | 0.734 | 0.227 | 0.077 | (Delfin et al. 2014) |
| Sumatra | 72 | 48 | 0.771 | 0.229 | 0.021 | (Gunnarsdottir et al. 2011) |
| Taiwan | 549 | 299 | 0.669 | 0.308 | 0.084 | (Ko et al. 2014) |
| Oceania | 1331 | 650 | 0.689 | 0.277 | 0.106 | (Duggan et al. 2014) |
| Siberia | 525 | 244 | 0.574 | 0.336 | 0.217 | (Duggan et al. 2013) |
| Mexico<sup>a</sup> | 113 | 90 | 0.867 | 0.133 | 0 | (Mizuno et al. 2014) |

Note: The proportions do not sum up to 1 since some haplotypes are shared both within and between populations. a. The individuals from Mexico are all of Amerindian origin.

Figure 4 shows the proportion of pairs of sequences shared between and within NWA populations. Siriano, Other-ET, and Pasto were the only groups without shared haplotypes within the populations. The majority of between-group haplotype sharing involved Arawakan and Eastern Tukanoan groups. The Arawakan groups share mostly with groups living in close proximity (Figure S4), e.g. Yucu-Matapi shared with Tanimuka; Curripaco with Puinave and Nukak, and Ach-Piapoco with Saliba and with the Guahiban groups Sikuani and Guayabero. In contrast, most Eastern Tukanoan groups, who practice linguistic exogamy, shared haplotypes among each other (except for Tanimuka, who shared only with Yucu-Matapi). In contrast, the Western Tukanoan groups Siona and Coreguaje shared primarily within their populations and did not share haplotypes with the Eastern Tukanoan groups.

The groups from the Andean foothills--Inga, Kamentsa, and Pasto--showed different patterns of shared haplotypes, despite the fact that they live in close geographic proximity. The Pasto, a group that has lost its native language and that is highly incorporated into the admixed local population, shared no haplotypes with any population. The Kamentsa shared haplotypes only with the Inga, while the Inga also shared haplotypes with three other groups located further inside the Amazonian area: Carijona, Coreguaje and Mur-Uitoto. Finally, of the three groups living on the banks of the Amazon River close to the town of Leticia, the Cocama shared with both the Yagua and Tikuna, whereas the latter two groups lacked common haplotypes.

**Figure 4.**
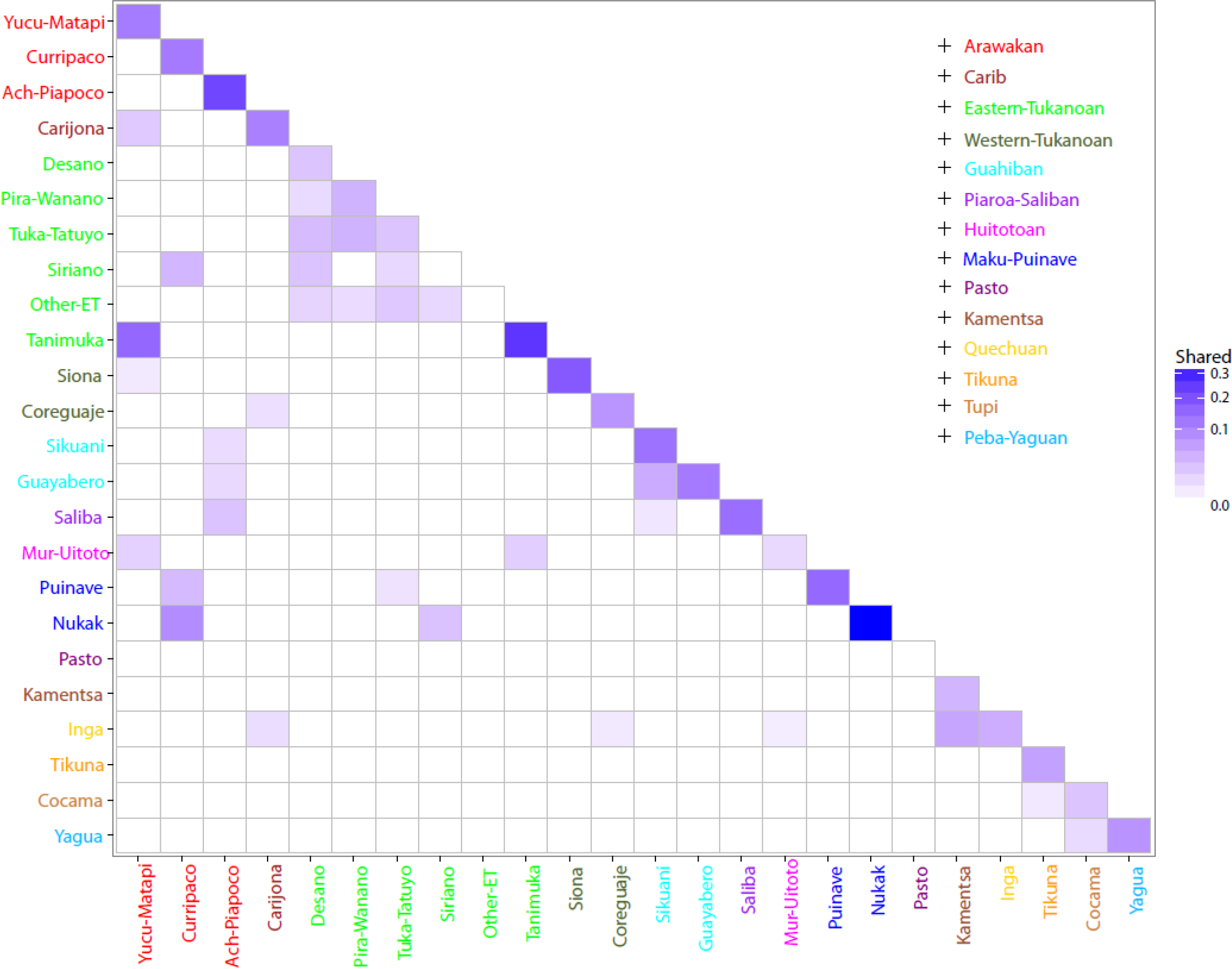
Matrix of shared haplotypes between populations. The color scale indicates the proportion of the total haplotypes that are shared within (on the diagonal) or between (below the diagonal) populations.

### Haplotype networks

The networks of haplotypes (Supporting information Figure S5 A-D) complement the patterns of sequence sharing, but in addition allow us to discern clusters of related (not just identical) haplotypes. We observed that some of these clusters were common among different language families and others were restricted to specific language families or to groups living in close geographic proximity; these are highlighted in Figures S5 (A-D). For instance, Arawakan and Eastern Tukanoan groups exhibited several haplotypes within haplogroups A2 (Cluster I, Figure S5A), B2 (Cluster I and II, Figure S5B), and C1 (Cluster III, IV, V and VI, Figure S5C) that were either shared or separated by only a few mutational steps. Notably, several of these clusters also included individuals speaking Maku-Puinave languages (cf. Cluster I and II, Figure S5B, and Cluster III, IV, and V, Figure S5C). Clusters of haplotypes restricted to specific groups are represented by clusters I and II in Figure S5D, exclusive to Eastern Tukanoan and Huitotoan populations, respectively. Furthermore, the haplotypes of the Inga (Quechuan) and the Kamentsa, who live in close proximity in the Andean foothills, were either shared between them or closely related (e.g. cluster II in Figure S5C). Finally, the haplotypes of the Guayabero and Sikuani (Guahiban) were mostly differentiated from those of other populations and generally shared by several individuals within the family (clusters II and III, Figure S5A; cluster I, Figure S5C). The sequences belonging to cluster I in haplogroup C (cluster I, Figure S5C) lack the diagnostic mutation A13263G for haplogroup C, but contain other diagnostic mutations that allow unambiguous assignment to haplogroup C. MtDNAs with this variant were previously identified in eastern Colombia by RFLP typing (Torres et al. 2006), where they occurred at high frequency in Guahibo, Piapoco, and Saliba groups. Given their high frequencies in the Guahiban groups, these haplotypes appear to belong to an autochthonous lineage that has then diffused into other groups living in the Orinoco basin.

### Genetic structure and genetic distances

The AMOVA analysis (Table 4) allows us to test different hypotheses concerning how genetic variation is structured in NWA. We defined groups *a priori* based on language affiliation, geographic proximity, and distribution along major rivers or their tributaries to evaluate how much of the observed variation is explained by each grouping strategy. We observed that of the three grouping strategies, grouping populations by their distribution along rivers resulted in the largest among-group component of the genetic variance. In contrast, language was a poor predictor of the genetic structure, showing negative and nonsignificant values for the component of variance due to differences among groups, indicating larger genetic differences among groups of populations speaking related languages than among linguistically different groups. Finally, geographic proximity was also a poor predictor; although the among-group component was larger than for language, it was not significantly different from zero. An important aspect to note is that although grouping by rivers performed better than grouping by geography or language, it still did not provide a very good description of the genetic structure, since the percentage of variance due to differences among populations within groups was still higher than the among groups component, suggesting the existence of other levels of substructure within populations.

**Table 4.**
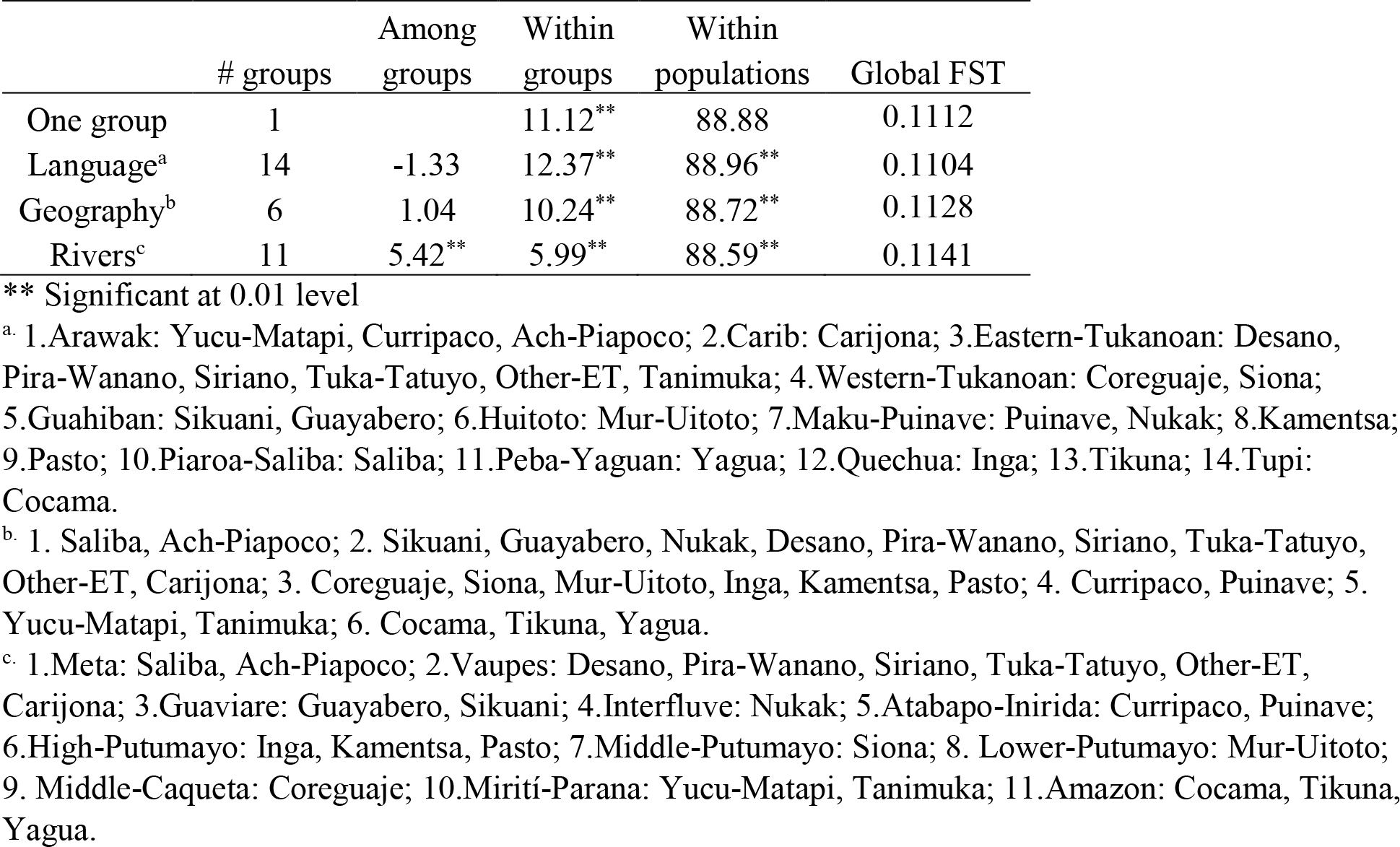
Analysis of molecular variance (AMOVA)

The matrix of genetic distances between populations based on pairwise Φ_ST_ values (Supporting information Figure S6) was used to construct an MDS plot (Figure 5). The populations do not form any clear clustering: the majority of populations are grouped together in the center of the plot (indicated by the inner circle in Figure 5) with an average pairwise Φ_ST_ = 0.03, while around the main cluster a second group of populations showed higher differentiation (external circle, average Φ_ST_ = 0.07). Finally, Sikuani, Siona, and the hunter-gatherer Nukak appeared as outliers with high genetic differentiation (average Φ_ST_ = 0.22). This picture did not change after adding an additional dimension to the MDS plot (Supporting information Figure S7). Particularly striking were the small genetic distances between the Eastern Tukanoan groups, who clustered together in the center of the MDS plot. Although Tanimuka appeared more distant from the main cluster of Eastern Tukanoan groups, their pairwise Φ_ST_ values were not significantly different (Supporting information Figure S6) and the average Φ_ST_ (0.02) indicated low genetic differentiation among all Eastern Tukanoan groups. In contrast, the Coreguaje and the Siona, who speak languages of the Western Tukanoan branch, showed larger genetic distances, both with the Eastern Tukanoan groups and with each other. Populations from each of the other language families did not form clusters with their linguistic relatives. For example, Arawakan groups occupied different positions in the plot and their Φ_ST_ values were significantly different.

**Figure 5.**
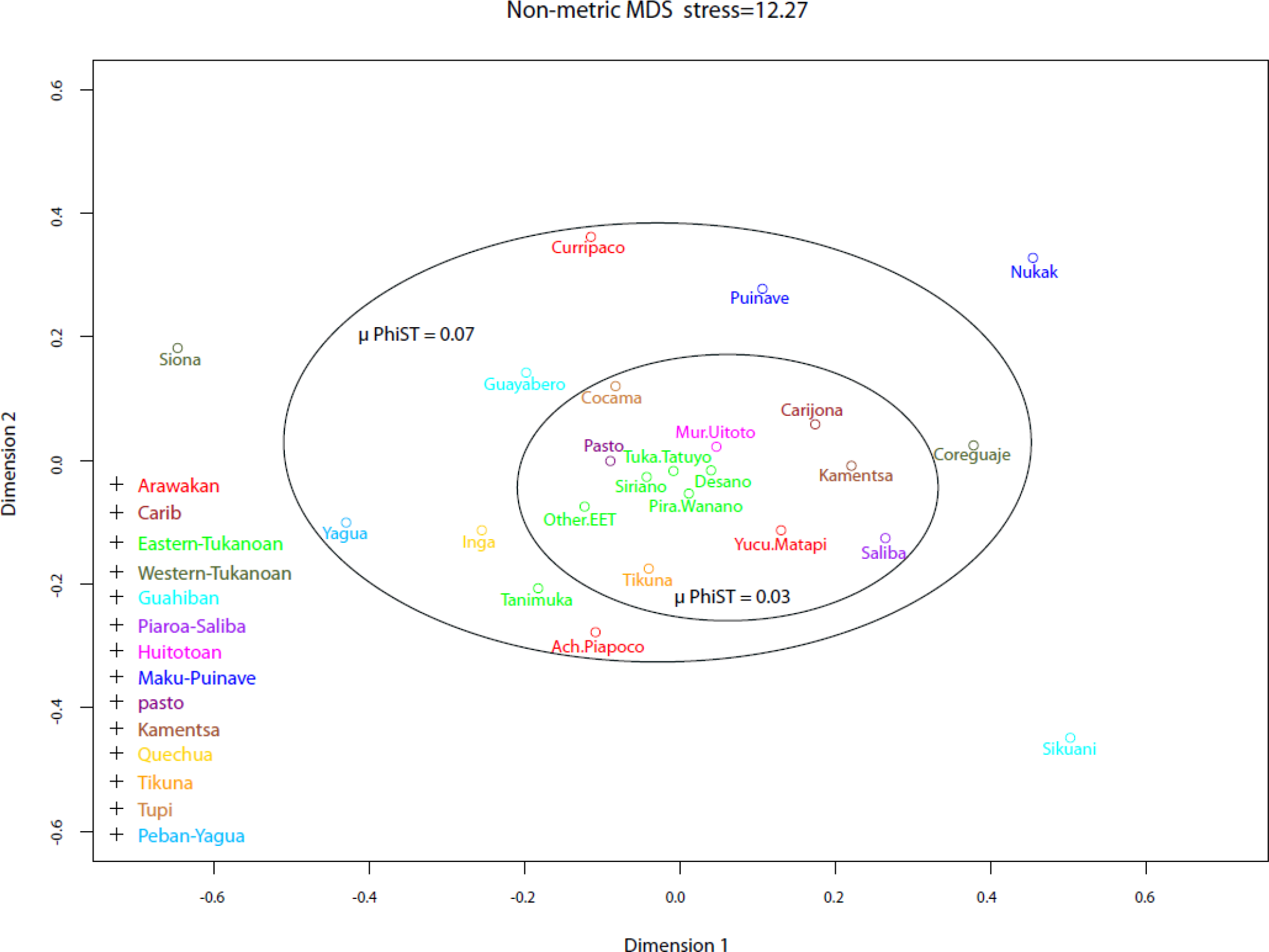
Multidimensional Scaling plot based on Φ_ST_ genetic distances. Stress value is given in percentage. The inner circle indicates populations with low genetic differentiation and the outer circle indicates populations with moderate differentiation. μ PhiST is the average pairwise Φ_ST_ value within each circle.

The results of the Mantel test showed a lack of significant correlation between geographic distances, estimated as great-circle distances, and the matrix of pairwise Φ_ST_ values (r = 0.07, p-value = 0.28). However, since rivers emerged as an important factor explaining the structure of genetic variation in the AMOVA results (Table 4), we also performed a multiple regression analysis on distance matrices (MRM), where we added rivers as an additional predictor variable. Adding rivers to the regression model resulted in an increase in the amount of variation explained by the model (Table 5), with rivers being a significant predictor (p-value = 0.01). We then jack-knifed over populations (Pugach et al. 2016; Ramachandran et al. 2005) and identified three populations as outliers: Sikuani, Siona, and Nukak, who appeared as outliers in the MDS plot as well (Figure 5). We performed the multiple regression analysis excluding the outliers; this resulted in an increase of 3.4 % in the R square value, a better correlation between genetic and geographic distances, and geography becoming a significant predictor factor (p-value < 0.05) (Table 5, supporting information Figure S8), although rivers were no longer a significant predictor of genetic subdivision.

**Table 5.**
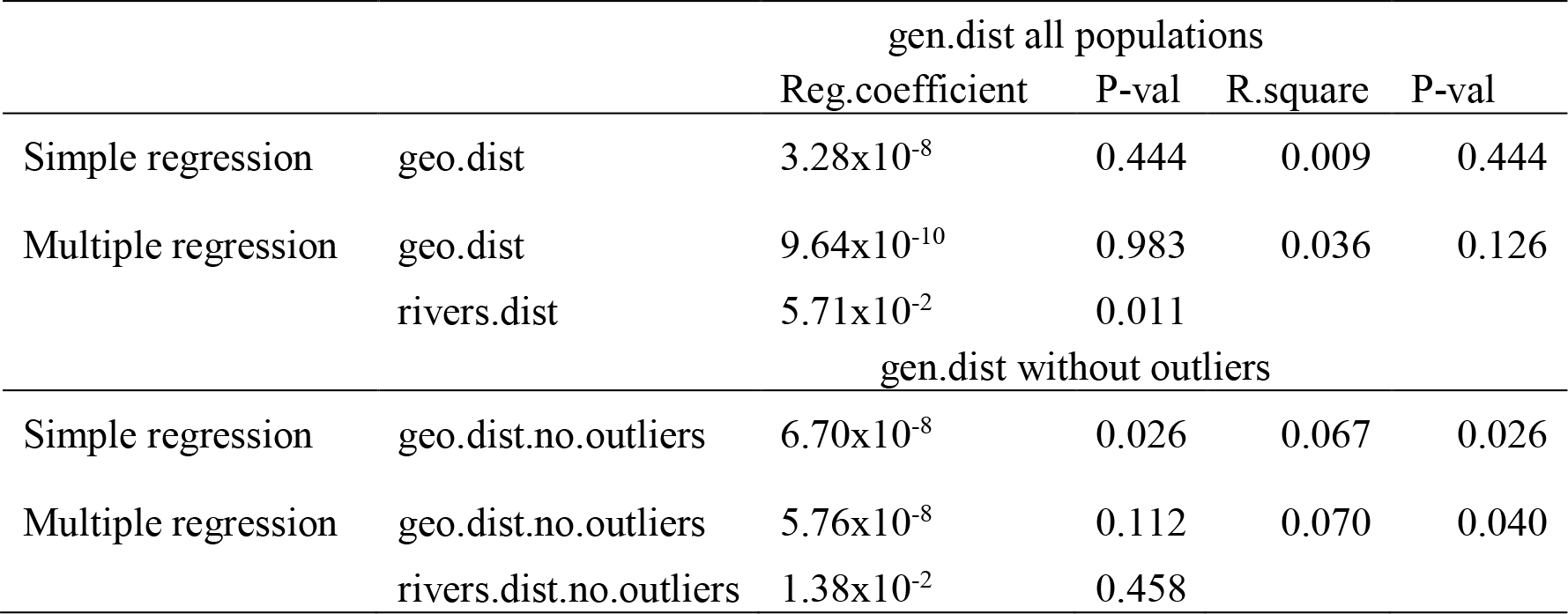
Multiple regression analysis on distance matrices

### Bayesian demographic reconstruction

Bayesian skyline plots (BSP) were generated by haplogroup (A2, B2, C1 and D1) and by population. All four haplogroups showed an increase in effective population size between 17,500 - 25,000 years before present. This signal was more evident for haplogroups A2 and C1, which have the highest number of sequences (Supporting information Figure S9). In contrast, the BSP plots by population showed different outcomes. We observed four main trajectories (Supporting information Figure S3): first, a signal of population size increase shown by Yucu-Matapi, Curripaco, Desano, Siriano, Inga, Pasto, Mur-Uitoto, Tikuna, and Cocama (exemplified by Yucu-Matapi in Figure S3A); second, population stability through time shown by Ach-Piapoco, Tanimuka, Coreguaje, Siona, Kamentsa, Puinave, and Yagua (exemplified by Coreguaje in Figure S3B); and third, population contraction shown by Sikuani, Guayabero, and Nukak (exemplified by Nukak in Figure S3C), which is particularly striking for Sikuani (Supporting information Figure S3D). These differences in the effective population size through time suggest that these populations have followed independent demographic histories.

## DISCUSSION

We have investigated the genetic diversity of ethnolinguistic groups from NWA at the level of complete mitochondrial genomes. This area is underrepresented in previous studies, and our data contribute to fill a gap in our knowledge about the genetic diversity of modern human populations. We have found that NWA harbors a considerable amount of genetic diversity, with evidence for contact among different ethnolinguistic groups, contrary to the common picture of Amazonian populations as small and isolated with low genetic diversity (Fuselli et al. 2003; Wang et al. 2007). NWA populations show values of nucleotide diversity as high as or higher than those observed in most other non-African populations (Supporting information Figure S10), and they display the second-highest amount of sequence sharing in a world-wide comparison (Table 3). The complete mitochondrial genome is the maximum level of resolution one can achieve to differentiate individuals and populations at the maternal level, so the presence of identical sequences among populations living in distant geographic areas indicates recent contact and/or common ancestry.

### Lack of genetic structure along linguistic lines

Although our dataset includes populations speaking languages belonging to different language families, we found that linguistic affiliation is a poor predictor of genetic structure, as shown by the AMOVA analysis (Table 4) and by the Correspondence Analysis based on sub-haplogroup frequencies (Figure 2). This indicates that language does not constitute a barrier to gene flow, and that groups have been interacting with other neighboring groups, especially along rivers, which in our analyses performed better in explaining the patterns of genetic diversity. Archaeological and linguistic evidence demonstrates that NWA has been an area of intense contact and movement of peoples of different cultural traditions, evidenced by the diffusion of ceramic styles (Heckenberger 2002; Lathrap 1970; Zucchi 2002) and shared subsistence strategies, by the existence of language areas and contact-induced linguistic change (Aikhenvald 1999), and the generalized multilingualism among groups (Sorensen 1967; Stenzel 2005). Likewise, cultural anthropology provides additional evidence of contact among groups. For example, both Arawakans and Eastern Tukanoans share a ceremonial complex for male initiation known as Yurupari, in which sacred flutes and trumpets are only played by males, as well as sharing myths concerning the hero Kúwai (Hugh-Jones 1979; Jackson 1983; Vidal 2002). In addition, the Eastern Tukanoan groups from the Pira-Parana and Apaporis rivers (Barasano, Makuna, and Tanimuka) reveal Arawakan influence, since they also practice dances with masks during the season of high abundance of the palm tree fruit pupunha *(Bactris gasipaes)* (Hugh-Jones 1979). The genetic distances among populations provide additional evidence in this regard: although the global Φ_ST_ value of 0.11 indicates moderate differentiation (Hartl and Clark 2007), this value is driven by three populations, namely the Siona, Sikuani, and Nukak. These are highly differentiated from the other populations, likely reflecting the effects of genetic drift due to bottlenecks, as indicated by the positive Tajima’s D values (Figure 3) and the distribution of pairwise differences (Supporting information Figure S2). When we exclude these populations, we observe an average pairwise Φ_ST_ of 0.07, and populations appear close together in the MDS plot (Figure 5), indicating low genetic differentiation among NWA populations.

In this general picture the Eastern Tukanoan groups stand apart, since they cluster together in the CA and MDS plots (Figure 2 and Figure 5), and their pairwise genetic distances are small and non-significant (Figure S6). Linguists have proposed a time depth for the Tukanoan family of 2000-2500 years, based on a comparison of the diversity in Tukanoan languages with the diversity in Romance and Germanic languages (Chacon 2014). The time depth of the Eastern Tukanoan branch (and thus the time to the most recent common ancestor of the Eastern Tukanoan languages) would be even more recent, which might indicate that the peoples speaking these languages share recent common genetic ancestry as well (at least on the maternal side). However, the Eastern Tukanoan groups practice linguistic exogamy, and the close genetic relationships among these populations might be the result of this marital system in which women move among different ethnolinguistic groups. The consequences of the linguistic exogamy are also evident in the gene diversity values and the patterns of shared haplotypes. Eastern Tukanoans are the groups with the highest values of gene diversity, and they share more haplotypes among themselves than with other non-Eastern Tukanoan groups. In addition, their haplotypes tend to be closely related, as can be seen in the phylogenetic networks (Supporting information Figure S5). Analyses of the Y-chromosome as well as nuclear markers will help to disentangle the effects of linguistic exogamy vs. recent common ancestry on the patterns of genetic variation among Eastern Tukanoan groups.

The Tanimuka stand apart from the other Eastern Tukanoan groups in the analyses, which may reflect their settlement further south, along the Apaporis and Mirití-Paraná rivers. Moreover, they do not participate in the exogamic system with other Eastern Tukanoan groups, but interact mainly with the Arawakan groups Yucuna and Matapi. This is reflected in the patterns of haplotype sharing (Figure 4) as well as in their language, which shows evidence of Arawakan influence (Barnes 1999; Chacon 2014).

### The role of rivers in structuring genetic variation

Besides language, geography is another important factor in structuring the patterns of genetic variation in human populations (Ramachandran et al. 2005; Schonberg et al. 2011; Wang et al. 2007). One of the most salient characteristics of the physical landscape of NWA is the high density of rivers that drain the area, and their importance for human populations was earlier recognized by explorers and ethnographers that traveled through the region (Koch-Grünberg 1995; Wallace 1853). We found that the distribution along rivers is an additional important factor influencing the genetic structure of NWA populations: our AMOVA analyses (Table 4) show that clustering populations according to the rivers where they are distributed explains more of the genetic variation that is due to differences among groups than does grouping them by linguistic affiliation, i.e. populations living on the same river basin or in closely connected rivers are genetically more similar than those living on different rivers. This pattern is also observed in the distribution of sub-haplogroups among populations (Figure S1). For example, Curripaco and Puinave, who live on the Inírida and Atabapo rivers, are located close together in the plot, and the presence of Coreguaje, Yucu-Matapi, and Mur-Uitoto in the center of the plot could reflect their presence in a region where the Putumayo and Caquetá rivers are separated by their shortest distance, therefore facilitating contact among people inhabiting the basins and tributaries of these two rivers. Indeed, one Murui individual was sampled in a Coreguaje community, and two Uitoto individuals were sampled in the Mirití-Parana region, thus providing evidence for the movement of people among these groups. The results of the MRM analysis provide additional evidence in this regard: even though no correlation between genetic distances and geographic distances was observed via the Mantel test, we observed that adding river distances as an additional predictor variable resulted in an increase of around 3% of the R-square value (Table 5), indicating that rivers contribute to explaining a slightly higher percentage of the variation observed in the genetic distances.

Rivers in Amazonia serve a double function in providing a means of communication as well as subsistence, and the wide distribution of certain cultural traits (e.g., the production of Saladoid-Barrancoid ceramics and circular plaza village settlement patterns), has been associated with the expansion of Arawakan-speaking populations along the extensive system of NWA waterways (Heckenberger 2002; Hornborg 2005; Lathrap 1970; Lowie 1948). They also mark a distinction in subsistence strategies between the more numerous “river people” that build canoes, settle along rivers, and rely on horticulture and fishing, and the “forest people” that inhabit the interfluvial areas, settle away from the major rivers, and base their subsistence on foraging (Epps and Stenzel 2013). Additionally, the rivers have profound meanings and are embedded in the cosmogonies of several NWA indigenous groups. The Eastern Tukanoan creation myths describe the journeys that the ancestors of the people made to settle this world on board an anaconda canoe that travelled up the Vaupes River; from the anaconda’s body all the Tukanoan siblings emerged (Chernela 2010; Jackson 1983). Arawakan groups also describe a series of ever returning voyages from the sacred center of the world and the place of emergence of the first ancestors at the rapids of Hípana on the Aiary River, covering the major arteries of the Rio Negro, Orinoco, and Amazon (Wright 2002; Zucchi 2002). Therefore, our findings about the role of rivers in structuring the genetic variation are in keeping with the pivotal role that rivers play for NWA populations.

The lack of fit between genetic and simple geographic distances may be the result of relatively recent movements and the displacement of ethnolinguistic groups from their traditional territories. Population dynamics and population sizes were drastically altered during the last five centuries, starting with early colonial times (16^th^ and 17^th^centuries), when many groups were decimated by newly introduced epidemics and moved away from the accessible margins of the major rivers to avoid the slave raids of the Spanish, Portuguese, and Dutch colonizers. Similar perturbations happened during the time of the Christian missions in the 18^th^century, when many groups were forced to relocate to multiethnic mission settlements, and finally during the rubber boom between the 19^th^ and beginning of the 20^th^centuries, when the groups who managed to escape the mercenaries exploiting the rubber fields resettled in remote areas in the headwaters of small rivers (Dixon and Aikhenvald 1999; Hill and Santos-Granero 2002; Stenzel 2005). The inferred reduction in population size of the Tanimuka, Sikuani, Guayabero, and Nukak, as indicated by their low diversity values, the positive Tajima’s D values (Figure 3), the distribution of pairwise differences (Figure S2), and the reconstruction of effective population sizes (Figure S3C, D), might be a result of these social upheavals.

### The impact of subsistence strategies on the genetic diversity

NWA contains groups with different subsistence strategies, with manioc (*Manihot esculenta*) as the main staple among horticulturist groups, who are best described as riverine horticultural societies, given their close association with rivers. The Nukak, in contrast, are traditionally foragers, who still rely on hunting and gathering and move throughout the extensive area between the Guaviare and Inírida rivers. Furthermore, the Guayabero, Sikuani, and Puinave are traditional foragers who have only recently undergone the transition to agriculture, and are therefore considered as HGP together with Nukak in our analyses (Table 1). Our data show that agricultural societies (AG) have higher levels of diversity on average than forager groups (HGP) as indicated by the Mann-Whitney U test (P-value = 0.03), while the HGP groups have larger values of Tajima’s D statistic (Figure 3) and do not show signals of population expansion (Figure S3). These findings agree with patterns reported for other hunter-gatherer populations around the world (Aime et al. 2013; Excoffier and Schneider 1999; Oota et al. 2005) and contrast with the genetic signature of an agricultural way of life, namely higher effective population size (Patin et al. 2014), higher levels of diversity, and significantly negative values of Tajima’s D test (Aime et al. 2013).

However, subsistence strategies are flexible and diverse among NWA populations. Horticulturalists complement their diet with occasional hunting and/or gathering of several kinds of palm fruit, and extensive exchanges between AG and HGP groups have been reported. In this system, HGP populations usually provide meat and several products from the forest, such as the poison curare for the tips of darts and arrows, in exchange for different cultivated products, such as manioc and other trade goods (Epps and Stenzel 2013; Jackson 1983; Milton 1984). Nonetheless, this exchange seems to be exclusively restricted to goods and labor, with little or no intermarriage documented between AG and HGP groups (Aikhenvald 1996). In contrast, we observed shared haplotypes between AG and HGP groups, which likely reflects intermarriage or recent common ancestry. For example, the most frequent haplotype in the Arawakan AG group Curripaco (Haplotype H_84 in Figure S4) is observed at high frequency in the HGP Nukak (and in the Eastern Tukanoan AG group Siriano). Moreover, the HGP Puinave share several haplotypes with the AG group Curripaco (H_219, H_161, H_117 in Figure S4), a likely result of intermarriage between these groups, since there are communities on the Inírida River where one finds individuals from both groups. Similarly, the Guahiban HGP groups Sikuani and Guayabero exhibit a haplotype at high frequency (H_43 in Figure S4) that is shared with the AG Ach-Piapoco as well as further haplotypes related to haplotypes found in AG Arawakan groups (cluster I Figure S5C and cluster II S5B). This may reflect contact among them, since there are Piapoco communities on the lower Guaviare River as well as Sikuani communities on the Meta River, places where these groups overlap. However, it is difficult to determine the direction of the gene flow or to distinguish between contact and common ancestry as explanations for shared mtDNA haplotypes. Nevertheless, it is plausible that where haplotypes are shared the source population is the one in which the haplotype is present at higher frequency. For instance, the shared haplotype between the HGP Puinave and the AG Curripaco (H_219 in Figure S4) has a likely origin in Puinave, because of its higher frequency and the presence of related haplotypes in Puinave (cluster I Figure S5B). The source of the shared haplotype among the HGP Nukak and the AG Curripaco and Siriano (H_84 in Figure S4) is more difficult to infer, since its frequency is similar in the Nukak and in the Curripaco; furthermore, three other haplotypes present in the HGP Nukak and Guayabero are only one mutation apart from it (cluster II Figure S5B). Therefore, it is likely that this haplotype, too, moved from the HGP populations into the AG Curripaco. A similar explanation could be given for H_43 in Figure S4, which is part of the cluster I in Figure S5C, moving from the HGP Guayabero and Sikuani into the AG Ach-Piapoco. Thus, these observations seem to fit a scenario of asymmetric gene flow in which women move from HGP to AG, a pattern that has been reported for populations in Central and Southern Africa (Barbieri et al. 2014; Destro-Bisol et al. 2004; Verdu et al. 2013). However, this scenario will be further refined by analyses of Y-chromosome and genome-wide data, which will allow us to determine whether the gene flow among groups was sex-biased (i.e. involving the movement of only females or only males among groups) and to make inferences about the time and magnitude of these events.

In conclusion, this study provides new data from this remote and little-studied part of the world, which allow insights into the impact of cultural practices on the patterns of genetic variation and on the population dynamics of NWA groups. Although our current data do not allow us to distinguish whether the population movements took place prior to European contact or only later, analyses of Y-chromosome variation and genome-wide data will shed further light on the genetic history of NWA. Furthermore, historical genetic studies will benefit from more archaeological work in NWA, since huge areas remain completely unexplored.

## Acknowledgments

We are grateful to all sample donors, communities, community leaders, and regional indigenous organizations. L.A. especially gives thanks to Consejo Regional Indígena del Guaviare, Asociación de Autoridades Tradicionales y Cabildos de los Pueblos Indígenas del Municipio de Leguízamo y Alto Resguardo Predio Putumayo, Asociación de Cabildos Indígenas del Trapecio Amazónico, Organizatión Zonal Indígena Del Putumayo, the staff of Parques Nacionales Naturales in Puerto Leguízamo, the office of Indigenous Affairs in Puerto Inírida, Rafael Rodriguez, William Yucuna, the late Gustavo Arias, and all people who helped during the fieldtrips for their valuable collaboration and warm welcome during our stay in their communities. We also acknowledge Roland Schroeder for laboratory technical assistance, and Enrico Macholdt, Alexander Huebner, Irina Pugach, and Michael Dannemann for advice with the data analyses. B.P. acknowledges the LABEX ASLAN (ANR-10-LABX-0081) of Université de Lyon for its financial support within the program “Investissements d’Avenir” (ANR-11-IDEX-0007) of the French government operated by the National Research Agency (ANR). L.A. was supported by a graduate grant from COLCIENCIAS; research was supported by funds from the Max Planck Society.

---

**Supporting information Table S1.**
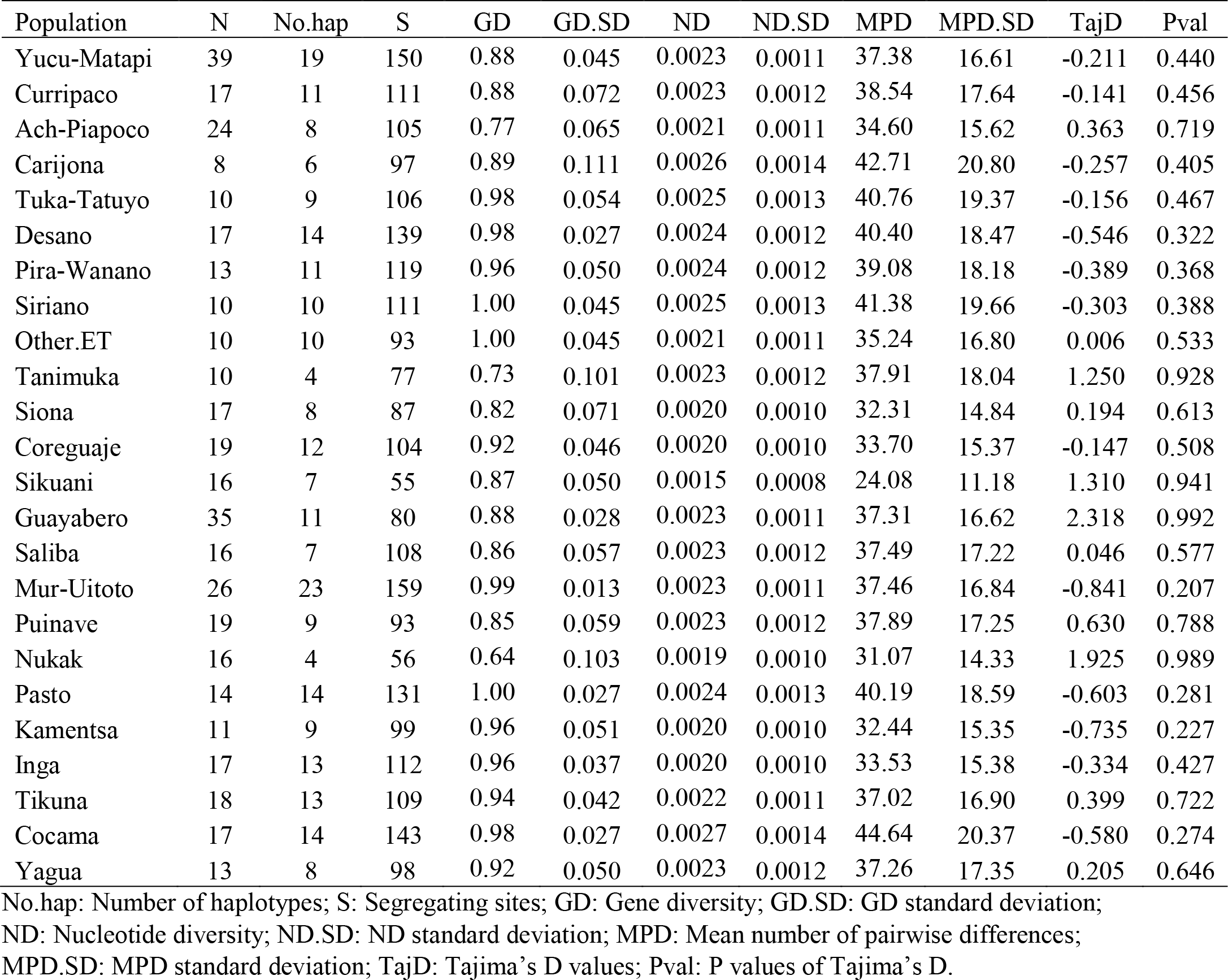
Molecular diversity indices by population

